# Meta-analysis of the effects of sleep deprivation on depression in patients and animals

**DOI:** 10.1101/2020.03.02.971689

**Authors:** Baiqi Hu, Mengfei Ye, Dan Feng, Jianghua Ying, Tingting Mou, Fangyi Luo, Tingting Lv, Liya Jiang, Chao Qian, Zhinan Ding, Chaoyang Yu, Hui Gao, Jian Zhang, Zheng Liu

**Author notes:** Corresponding author at: Department of Behavioral Neurosciences, Science Research Center of Medical School, Shaoxing University, Shaoxing, Zhejiang, China, 312000. These authors contributed equally to this work (Baiqi Hu, Mengfei Ye, Dan Feng).

## Abstract

**Objective:** Research on the antidepressant effects of sleep deprivation (SD) is lagging and has not produced completely uniform results in humans and animals. The present study aimed to reassess the effect of SD on patients and animals by meta-analysis based on updated research.

**Methods:** We searched PubMed, Embase and Cochrane Library for articles since the first relevant literature published up to June 10th, 2019. Data on sample characteristics, features of SD, and tests for depression were extracted.

**Results:** Fourteen articles were included, eight on humans and six on animals. We found that when the duration of SD in patients was 7–14 days, it reflected antidepression [-1.52 (−2.07, −0.97); I^2^=19.6%]. In animals, the results of sucrose consumption experiments showed that SD has depressogenic effects [-1.06 (−1.63, −0.49); I^2^=81.1%], while the results of forced swimming experiments showed that SD treated depression [-1.17 (−2.19, −0.16); I^2^=80.1%], regardless of the duration of sleep deprivation.

**Conclusion:** SD can be an effective antidepressant measure when the duration is 7–14 days in patients. In animal studies, SD has shown more antidepressant effects when measured by forced swimming experiments, whereas using sucrose consumption tests had the effect of worsening depression.

## 1. Introduction

Depression is a common, debilitating, and potentially lethal disorder that can affect people of all ages [1]. Over 300 million people worldwide suffer from depression; the World Health Organization (WHO) ranks it as the single largest contributor to global disability, accounting for 13.4% of “years of life lived with a disability” in women and 8.3% in men [2, 3]. Close to 800 000 depression patients die due to suicide every year. Suicide is the second leading cause of death in 15-29-year-olds [4]. Since relapse rates for depressive disorder are high, various potentially negative long-term outcomes are associated with it, including difficulties with interpersonal relationships, efficacy, tolerability and acceptability of antidepressants [5, 6]. Most people with depression have tried at least one antidepressant medication, although medication effects are slow to manifest, and side effects such as insomnia and anxiety lead patients to try different medications or refuse medication altogether [7, 8]. Furthermore, 30%–40% of patients are resistant to available antidepressant medications commonly prescribed for the major depressive disorder [9].

As a result of difficulties encountered when treating depression, there is an urgent need to find a nonpharmacologic therapy for it. In clinical practice, many nonpharmacologic therapies have attracted special attention, such as sleep deprivation (SD)[7], bright light therapy (BLT) [10], cognitive behavioral treatment (CBT)[11], and repetitive transcranial magnetic stimulation (rTMS)[7]. Among these, sleep deprivation therapy is one of the most rapid antidepressant interventions known [12]. Some clinical studies have shown that sleep deprivation (SD) is an effective treatment for patients with depression [13, 14]. Total sleep deprivation (TSD) for one whole night was found to improve depression symptoms in 40%–60% of patients [15]. Unfortunately, the therapeutic effects of SD are transient, and the depression symptoms can even return after a subsequent full night of sleep [7, 16]. Some results have indicated that patients who use a combination of antidepressants and SD have a significantly lower tendency to relapse after a full night’s sleep than those who do not [17]. Therefore, we hypothesize that some combinations of depression therapy can enhance therapeutic effects of SD.

In the present study, we aimed to explore the effectiveness of SD on depression. The antidepressant effects of SD have often been reported in humans, yet despite a recent meta-analysis [7], comprehensive aggregated data are lagging. Literature on SD lacks randomized controlled trials and has shown inconsistent results. The literature is not up-to-date, as the most recent study on SD was published in 2009. The duration of sleep deprivation has not been standardized across studies, which may have led to inconsistent results, so we explored whether SD treatment for patients with depression requires a more specific treatment course. In animals, the effects of SD have not been completely uniform. Animal models are a cornerstone of human research, particularly research on depression at the level of tissues, cells, molecules, and genes. However, no relevant meta-analyses have provided comprehensive results regarding animals. This article, using meta-analyses, provides an update on the effects of SD on patients and explores the effects of experimental SD on animals. At the same time, we discuss and evaluate whether sleep deprivation has a consistent effect on depression in animals and humans.

## 2. Methods

### 2.1 Literature search strategy

Studies related to the effects of SD on depression in patients or animals were identified by searching three different electronic databases (PubMed, Embase and Cochrane Library) for articles since the first relevant literature published up to June 10th, 2019, using the keywords (“sleep deprivation” OR “sleep curtailment” OR “sleep restriction” OR “sleep loss”) AND (“depression” OR “mood disorders”) in the title/abstract. A total of 1164 records meeting both search terms were returned. We excluded unmatched studies by keyword (case, review, report, and meta-analysis) and then selected studies to include or exclude according to titles and summaries. Additionally, relevant original studies cited in the selected articles were also eligible for inclusion. Final inclusion was determined by reading the full text of the studies.

### 2.2 Inclusion criteria

All included studies in this article met the criteria described by the participants, intervention, comparison, outcome, and study design (PICOS) according to recommendations by PRISMA and supplemented with criteria by the Quality Assessment of Diagnostic Accuracy 2 and the Newcastle-Ottawa Scale.

#### Patients

Patients included were between the ages of 12–80 years who had been diagnosed with depression based on the Diagnostic and Statistical Manual of Mental Disorders (DSM) and International Classification of Diseases (ICD) criteria, regardless of depression type (bipolar or unipolar) and gender (P); sleep deprivation (I); comparison to control conditions, there was SD design in the experimental conditions (C); outcome measures of the Hamilton depression scale (HAMD), Beck Depression Inventory (BDI), and the Montgomery Asberg Rating (MADRS) (O); and RCTs (S). In addition, patients who had serious organic diseases or mental and somatic comorbidities and pregnant women were excluded.

#### Animals

Differing from the requirements for depressed patients, it was not necessary to establish depressive-like behavior models in animals before the intervention (P); experimental SD (I); comparison to control conditions, there was SD design in the experimental conditions (C); outcome measures of open field experiments, sucrose consumption tests, and forced swimming tests (O); and RCTs (S).

Articles lacking either the full text or primary data findings that could not be resolved with engauge digitizer were excluded.

### 2.3 Data extraction and quality assessment

Each article was read in its entirety by two researchers to extract the data and record the trial details in a standardized table containing the following information: author(s), year of publication, country, participant characteristics (e.g., sample size, age, gender, and sample type), SD characteristics (e.g., type and duration), adjunctive method (e.g., bright light therapy, cognitive behavioral treatment, and antidepressant drug), and outcomes for patients. Regarding animals, species, SD method, and depression test were also added. When no specific data were included—only graphs or figures—the authors were contacted and asked to provide the results of their experiments or the raw data. If that failed, data were estimated based on graphs or figures using a digital ruler[18, 19]. Primary data were estimated according to coordinate positions, and then statistical methods were used to calculate mean and SD. The risk of bias was estimated independently by two researchers (J. Y. and T. M.), who extracted and appraised the data, using the Cochrane Risk of Bias tool [20]. Inconsistencies between the two researchers were resolved through negotiation; when that failed, a third person was asked to judge.

### 2.4 Data synthesis and analysis

First, to assess the effects of SD on depression, we conducted a comprehensive analysis of the selected trials. Then, we performed a hierarchical analysis based on a significant variable(duration of SD)on patients. Subsequently, subgroup analysis was used to determine the sources of heterogeneity. We performed subgroup analysis by country and adjunctive method for the patient studies and by depression test for the animal studies. For each comparison, we numerated the standardized mean difference based on Hedges’ *g* as a measure of effect size, with value ranges of small (0.2–0.5), medium (0.5–0.8), and large (0.8 and above), as per standard convention. This approach can ignore differences in depression measurement tools so the analysis can be unified. We used the random-effects model by DerSimonian and Laird[21]. Funnel plots and the Egger test were used to examine the risk of effect size for small studies.

The heterogeneity of effect size within each comparison was tested using Cochran’s Q test and I^2^ statistics. Data were presented as effect size ± confidence intervals at 95%. Results were considered significant when the confidence interval range was lower or higher than zero and associated with a Cochran’s Q *p*-value lower than 0.05. All calculations were performed using Stata version 13.1.

## 3. Results

### 3.1 Study characteristics

Our search strategy resulted in 1164 articles from PubMed and other databases (Fig. 1). Redundant literature was eliminated, and literature was filtered for relevance according to keywords (case, review, meta-analysis, and report), after which 77 articles were excluded. After the removal of titles and abstracts, there were 30 articles that were screened by reading the full text. After excluding articles that lacked control group or primary data, a total of 14 studies meeting the inclusion criteria were ultimately included in our meta-analysis.

**Fig. 1.**
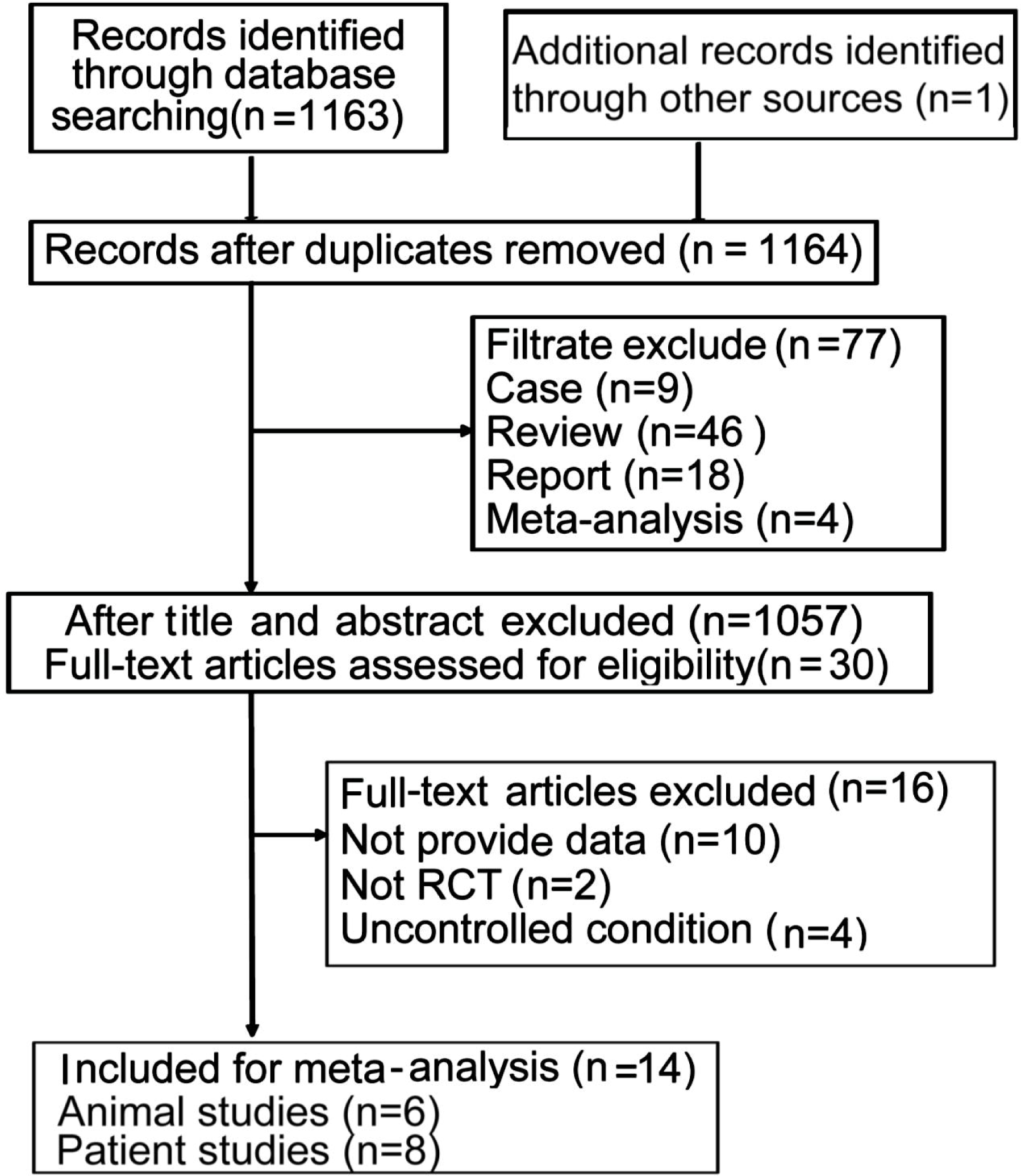
Selection process for trials included in the meta-analyses.

Among these, six were animal studies involving 13 trials and eight were patient studies involving 9 trials. For patient studies, TSD was applied in five articles, while partial sleep deprivation (PSD) was applied in three articles. No record of sleep curtailment, sleep restriction, or sleep loss was included. Most studies were conducted in Germany, the United States of America (USA), Turkey, Switzerland, and the Netherlands. All studies involved a combination of SD and other interventions. For instance, in human studies, six involved antidepressant drugs, and the other two involved, separately, BLT and CBT. Two of the animal studies were conducted on mice, and four were conducted on rats, including various species such as BALB/c, C57BL strains, Wistar, and Sprague-Dawley strains. The depression tests for animals included sucrose consumption tests, open-field tests, and forced swimming tests (Tables 1 and 2).

**Table 1.** Characteristics of the included patient studies.

| Author(s) | Country | Intervention | Sample sizes | Age (mean, year) | Type of depression | SM | Combined with other interventions | Duration | Depression scale |
| --- | --- | --- | --- | --- | --- | --- | --- | --- | --- |
| Elsenga et al., 1983[45] | Nether-lands | Clomipramine/SD<br>Clomipramine | 10<br>10 | 49.1±13.6<br>55.6±13.2 | Unipolar | TSD | Clomipramine | 7 days | Hamilton interview ratings |
| Trachsler et al., 1994[25] | Switzer-land | Trimipramine/SD<br>Trimipramine | 14<br>14 | 50.43±7.3<br>50.64±8.50 | Bipolar | PSD | Trimipramine | 28 days | HRS, MADRS |
| Kuhs et al., 1996[35] | Germany | Amitriptyline/LSD<br>Amitriptyline | 27<br>24 | 43.3±13.6<br>46.0±11.3 | Bipolar | LPSD | Amitriptyline | 14 days | HAM-D, 10 Item |
| Caliyurt et al., 2005[23] | Turkey | LPSD/Sertraline<br>Sertraline | 13<br>11 | 38.46±12.03 | Unipolar | LPSD | Sertraline | 14 days | HAM-D, 21 Item |
| Kundermann<br>et al., 2008[46] | Germany | TSD+CBT | 9 | 37±2.7 | Unipolar | TSD | CBT | 21 days | HDRS |
|  |  | CBT | 10 | 37.4±2.6 |  |  |  |  |  |
| Gorgulu et al.,<br>2009[24] | Turkey | TSD/Sertraline | 19 | 40±11.69 | Unipolar | TSD | Sertraline | 7 days | HAM-D |
|  |  | Sertraline | 22 | 33.27±11.18 |  |  |  |  |  |
| Smith et al.,<br>2009[47] | USA | TSD/Paroxetine | 7 | 69.0±4.6 | Unipolar | TSD | Paroxetine | 36 h | HDS-13 |
|  |  | TSD/Placebo | 6 | 68.6±4.9 |  |  |  |  |  |
|  |  | Paroxetine | 3 | 71.4±6.0 |  |  |  |  |  |
| Gest et al.,<br>2016[48] | Germany | Wake/BLT | 25 | 16.2±1.3 | Unipolar | TSD | BLT | one night | BDI-II |
|  |  | BLT | 37 | 15.8±1 |  |  |  |  |  |

Abbreviations: BDI-II = Beck Depression Inventory-II; BLT = Bright Light Therapy; CBT = Cognitive behavioral therapy; HAM-D = Hamilton
Depression Scale ;HAM-D, 10 Item = Hamilton Depression Scale (10-item) ;HAM-D, 21 Item = Hamilton Depression Scale (21-item); HDRS =
Hamilton Depression Rating Scale ; HDS-13 = Hamilton Depression Scale (13-item); HRS = Hamilton Rating Scale for Depression; LPSD = the
late partial sleep deprivation; LPSD = the late partial sleep deprivation; MADRS = the Montgomery Asberg Rating ; PSD = partial sleep
deprivation; SD = sleep deprivation; SM = sleep manipulation; TSD = total sleep deprivation; USA = United States of America

**Table 2.** Characteristics of the included animal studies.

| Author(s) | Country | Intervention | Sample sizes | Species | Strain | Gender | Age | Depression model | SM | Procedure | Duration | Depression tests |
| --- | --- | --- | --- | --- | --- | --- | --- | --- | --- | --- | --- | --- |
| Prathiba et al., 2000[42] | India | RSD<br>Control | 8<br>8 | Rats | Wistar | Male | 3 months | Clomipramine | REM SD | Pedestal above water | 4 days | OFT |
| Oliveira et al., 2004[49] | Brazil | Control<br>VEN-1 group<br>VEN-10<br>REMSD<br>REMSD+VEN-1<br>REMSD+VEN-10 | 20<br>20<br>20<br>20<br>20<br>20 | Rats | Wistar | Female | 3 months | Not specified | REM SD | Multiple-platform | 72h | OFT |
| Zhu et al., 2005[41] | China | REMSD<br>Control | 8<br>8 | Rats | Sprague Dawley | Male | Adult | CMUS | REM SD | Plowerpot | 72h | OFT |
| Jiang et al.,<br>2015[43] | China | RSD<br>Control | 10<br>10 | Rats | Sprague<br>Dawley | Male | 3 months | CMUS for 3<br>weeks | Not<br>specified | Platform above<br>water | 48h | SCT<br>OFT |
| Gonzalez et<br>al.,<br>2016[50] | Mexico | Males Control<br>Males 48-h PSD<br>Males 96-h PSD<br>Females Control<br>Females 48-h PSD<br>Females 96-h PSD | 12<br>12<br>12<br>12<br>12<br>12 | Mice | BALB/c | Both | 2 months | Not specified | Paradoxical<br>SD | Multiple-platform | 48h<br>96h | SCT<br>FST |
| Wang et al.,<br>2017[26] | China | SD for 3 days | 10 | Mice | C57BL/<br>6J | Male | 2-3<br>months | Not specified | Paradoxical<br>SD | Multiple-platform | 3 days<br>5 days | SCT<br>OFT<br>FST |
|  |  | Control | 10 |  |  |  |  |  |  |  |  |  |
|  |  | SD for 5 days | 18 |  |  |  |  |  |  |  |  |  |
|  |  | Control | 18 |  |  |  |  |  |  |  |  |  |
|  |  | FP5d+Sa | 10 |  |  |  |  |  |  |  |  |  |
|  |  | Control+Sa | 10 |  |  |  |  |  |  |  |  |  |

Abbreviations: CMUS = chronic mild unpredictable stress ;FST = forced swimming test; SCT = sucrose consumption test; OFT = open-field test;
P5d = paradoxical sleep deprivation for 5 Days; PSD = partial sleep deprivation; RSD/REM SD = rapid eye movement sleep deprivation; Sa =
saline; SD = sleep deprivation; SM = sleep manipulation; VEN-1 group = Venlafaxine-1 group

### 3.2 Study quality

#### Patients

Most studies adopted RCT, most random sequence generation indicated a low risk of bias[22]. Performance bias was not mentioned in most of the articles and was therefore mostly an unclear bias risk. Although the articles did not mention detection bias, the degree of depression was quantitatively measured by the depression scale; therefore, the tester factor had little influence, and the authors believed there was a low risk of detection bias. Two studies clearly did not blind participants and therefore had a high risk of bias, which can be considered the shortcomings of those studies [23, 24]. The final data for one study were unclear; thus, a high risk of bias was identified for the outcome of that study [25]. Unclear bias accounted for the majority of other biases since some literature only provided images instead of concrete data; thus the data obtained through software processing could have had some impact (Fig. 2A).

**Fig. 2.**
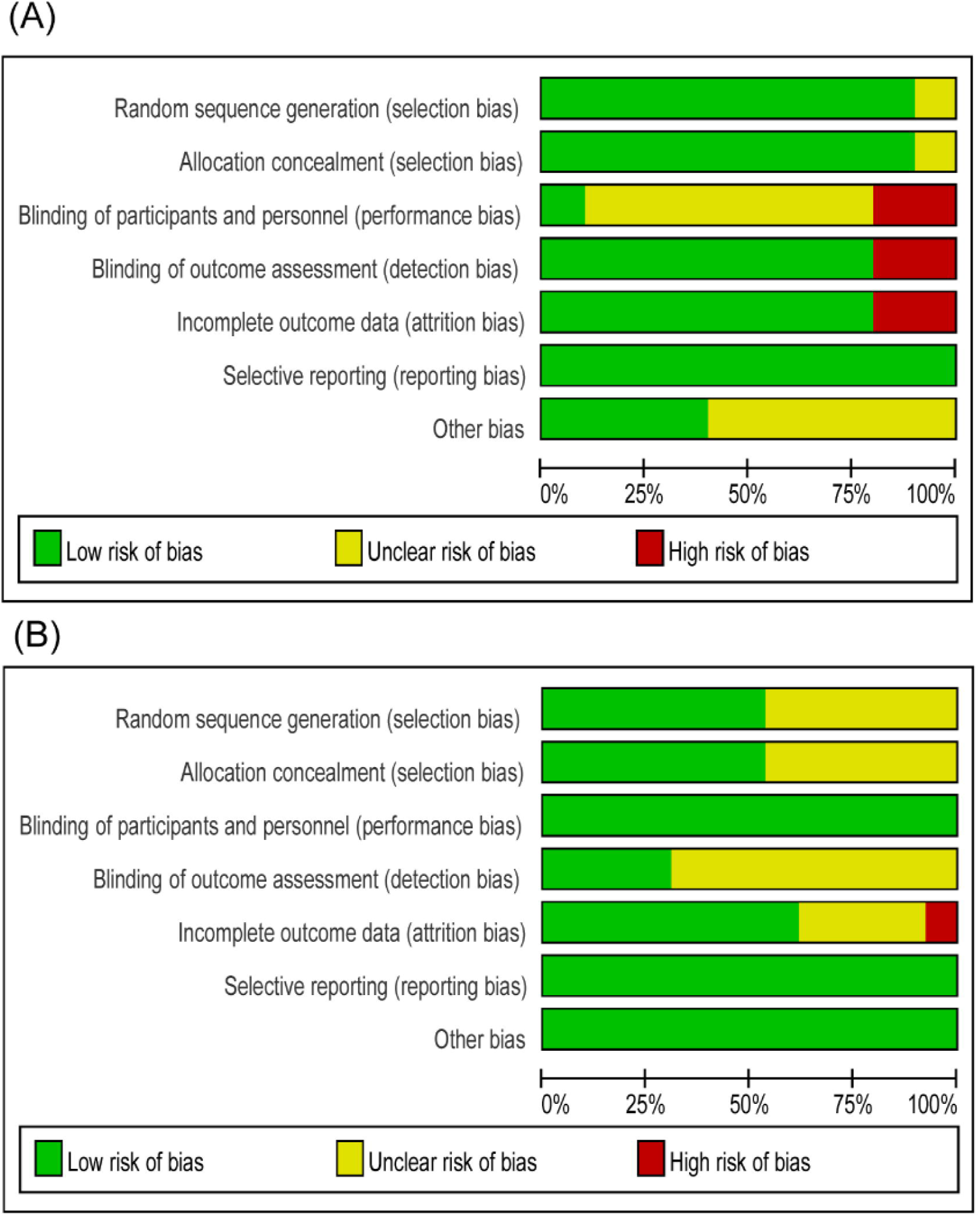
Risk-of-bias assessments of the included studies (domains from the Cochrane Handbook for Systematic Reviews of Interventions). A. Risk of bias assessments of the included patient studies. B. Risk of bias assessments of the included animal studies.

#### Animals

Participant blindness was not always mentioned, but in animal experiments, it was assumed to involve a low risk of bias. Only a few articles described the blinding method for study outcomes, which was considered as involving a low risk of bias, while the others were considered as having unclear bias without reference. Presentation of the results was complete in most articles, but one article did not provide the final data[26]; therefore, a high risk of bias was identified for the outcome of that study (Fig. 2B).

### 3.3 Main efficacy of the meta-analysis

#### Patients

Fig. 3 shows the total effect of SD on depression. Nine trials (10 datasets) reported depression using the HAMD. The random-effects meta-analysis elicited a summary effect size of −0.15 (95% CI, −0.80 to 0.50; I^2^=84.3%; P**<**0.001). When analyzed according to the SD schedule (**<**7 days, 7–14 days, >14 days), the forest plot showed that an SD duration of less than 7 days had a small effect of worsening depression [0.24 (−0.21, 0.69); I^2^=0%; P=0.43], a duration of 7–14 days had an antidepressant effect [-1.52 (−2.07, −0.97); I^2^=19.6%; P=0.288], and a duration of more than 14 days had the effect of worsening depression [0.76 (0.12, 1.40); I^2^=43.7%; P=0.169] (Fig. 4A).

**Fig. 3.**
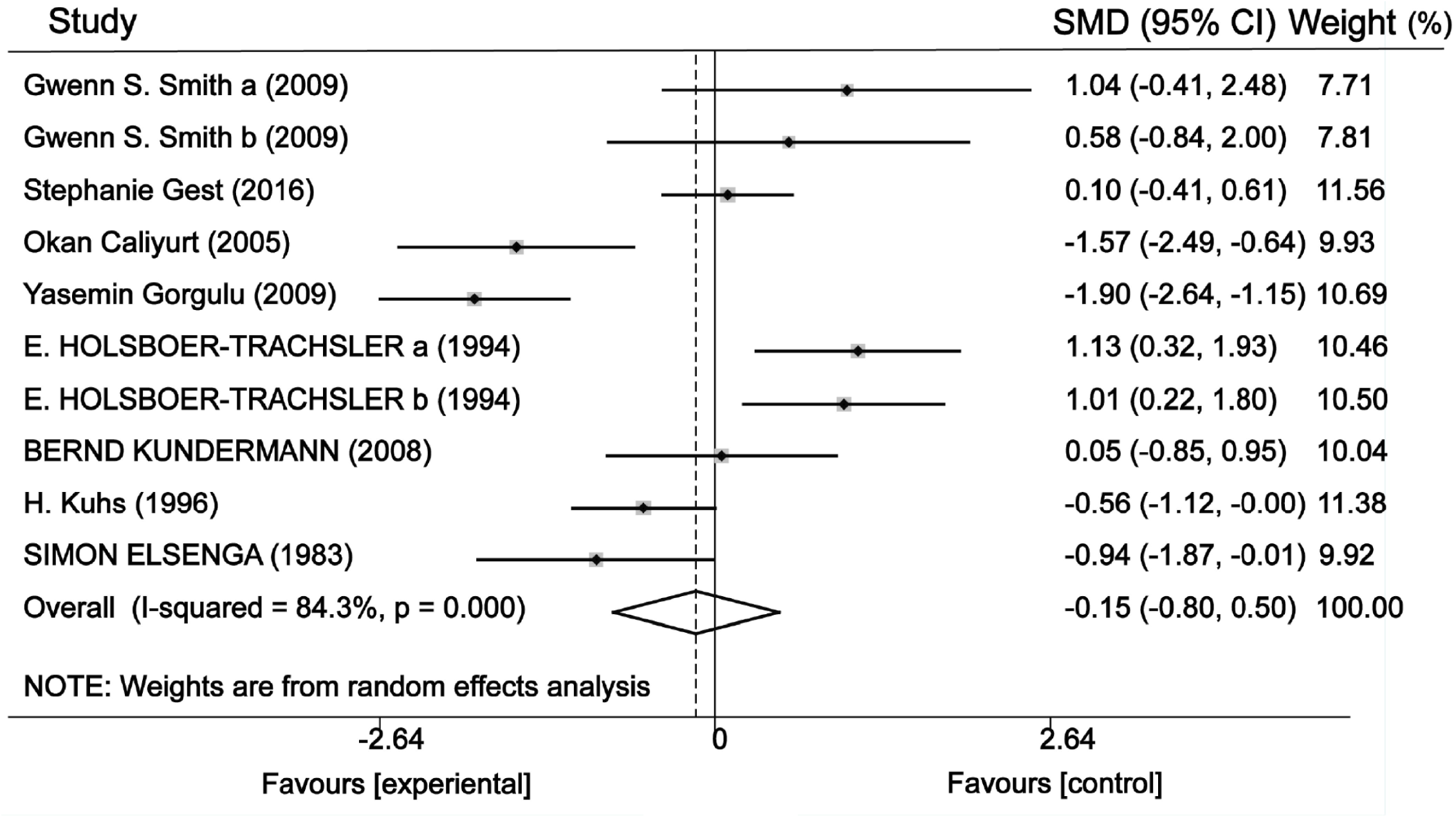
Forest plot of the random-effects model meta-analysis of the effect of SD on patients.

**Fig. 4.**
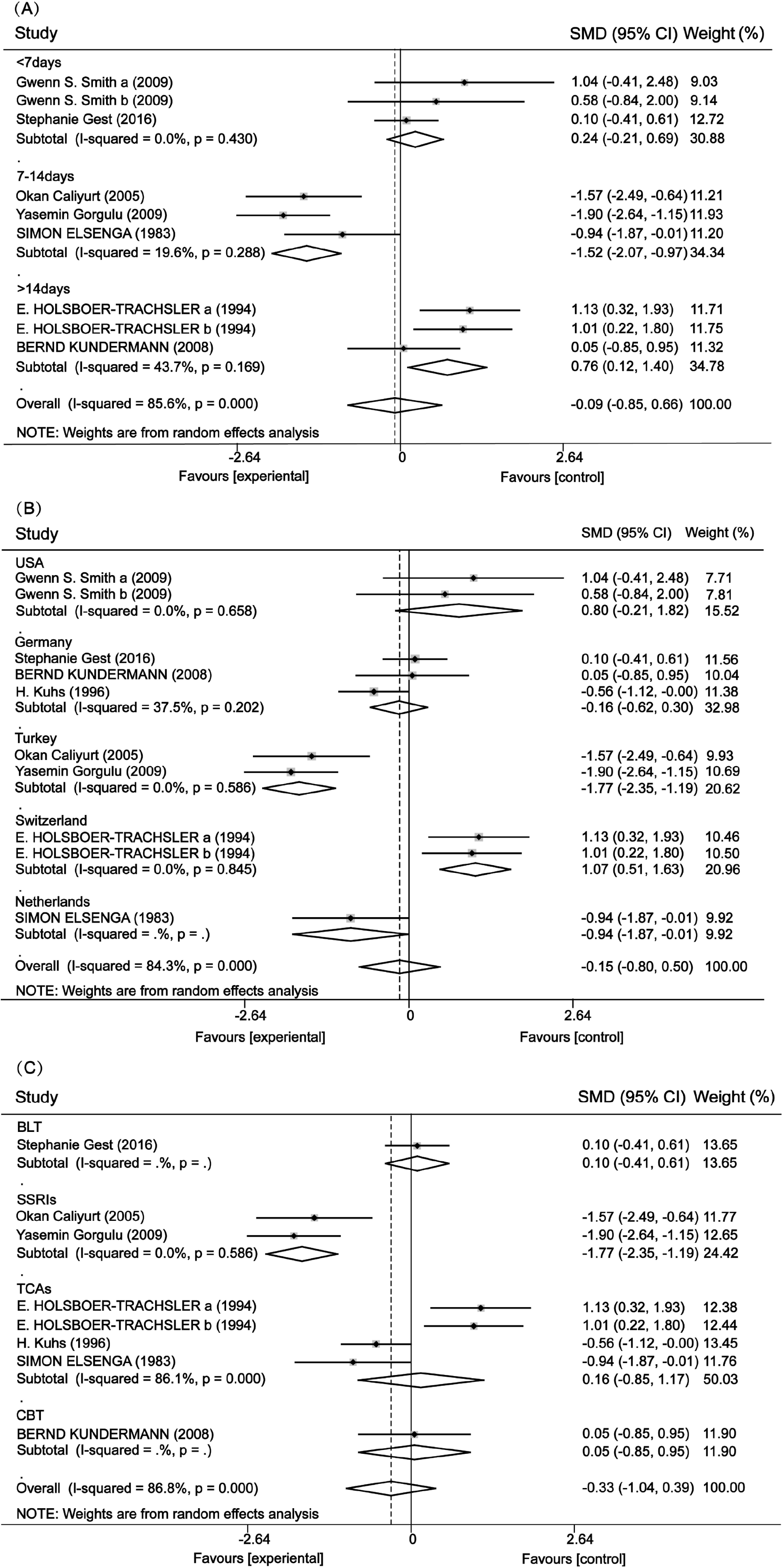
Forest plot of the random-effects model subgroup analysis of patients. A. Forest plot of the random-effects model subgroup analysis of patients with regard to the duration of SD. B. Forest plot of the random-effects model subgroup analysis of patients with regard to the country. C. Forest plot of the random-effects model subgroup analysis of patients with regard to combined therapy. Abbreviations: BLT = bright light therapy; CBT = cognitive behavior therapy; SSRIs = selective serotonin reuptake inhibitors; SD = sleep deprivation; TCAs = tricyclic antidepressive agents; and USA = United States of America.

#### Animals

The overall data suggested that SD had no significant effect on depression, and there was high heterogeneity [-0.28 (−0.73, 0.17); I^2^=86.8%; P**<**0.001] (Fig. 5).

**Fig. 5.**
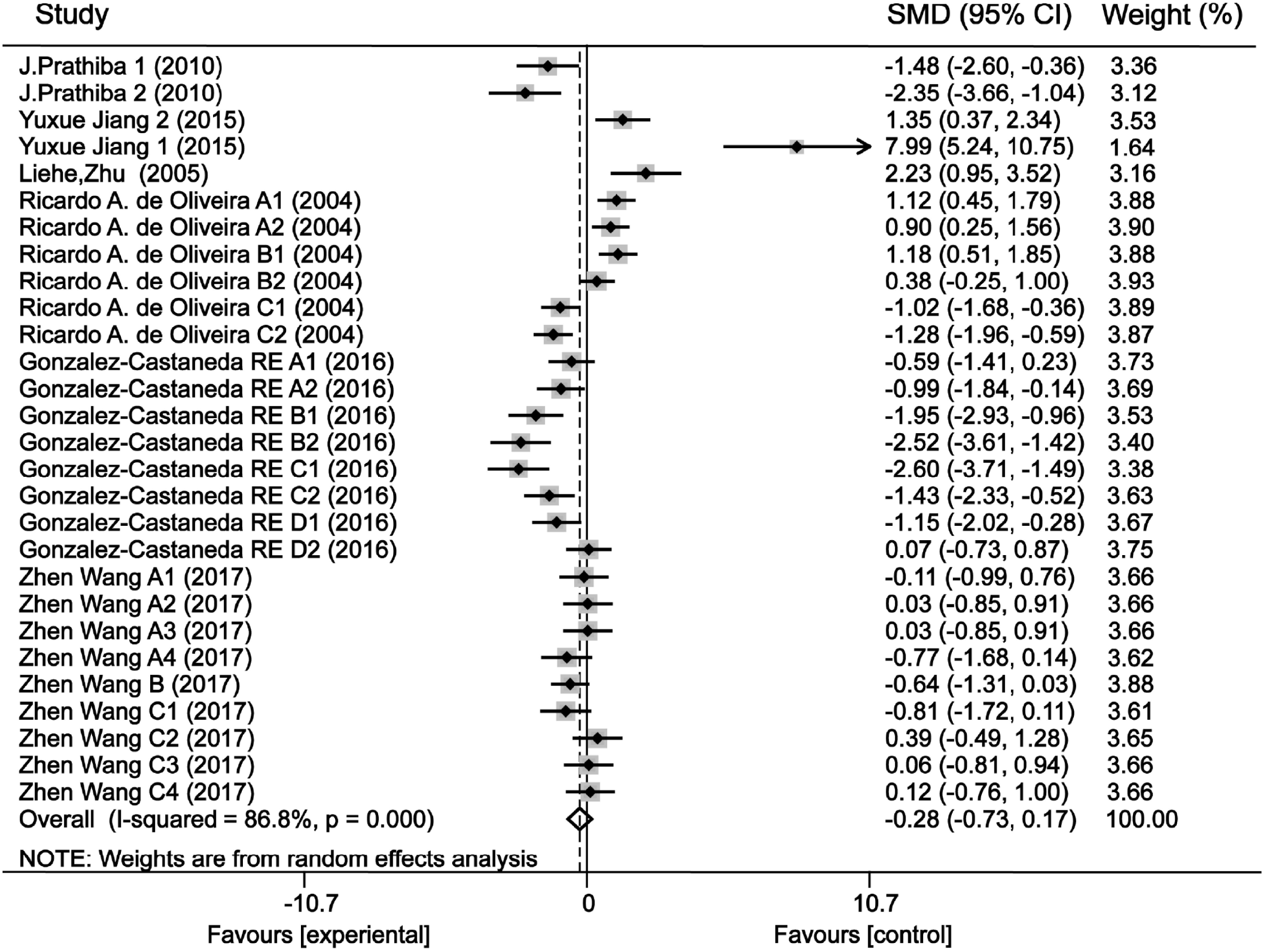
Forest plot of the random-effects model meta-analysis of the effect of SD on animals.

### 3.4 Heterogeneity analyses

Through subgroup analysis, we identified sources of research heterogeneity, which could have been related to the country where the research was conducted, the type of combined therapy employed, and the depression test that was used. For patients, the studies were divided into five subgroups according to country (Fig. 4B). Studies from Turkey showed high antidepressant effect sizes [-1.77 (−2.35, −1.19); I^2^=0%; P=0.586], while studies from Switzerland showed high effect sizes for worsened depression [1.07 (0.51, 1.63); I^2^=0%; P=0.845]. The studies were also divided into four subgroups for combined therapy (Fig. 4C). Studies that combined selective serotonin reuptake inhibitors (SSRIs) with SD showed an antidepressant effect [-1.77 (−2.35, −1.19); I^2^=0%; P=0.586].

The animal studies were divided into three subgroups according to the depression test used. Those using the sucrose consumption test to assess the level of depression indicated that SD worsened depression [-1.06 (−1.63, −0.49) (Fig. 6A); I^2^=81.1%; P**<**0.001], while those using forced swimming tests showed high antidepressant effects with SD [-1.17 (−2.19, −0.16); I^2^=80.1%; P=0.002] (Fig. 6B). Open-field tests showed no statistically significant differences [0.24 (−0.45, 0.92); I^2^=89.1%; P**<**0.001] (Fig. 6C). Sensitivity analysis revealed that the heterogeneity of the 7–14-day group decreased from 66.6% to 19.6%, indicating that the effect of SD on depression was related to its duration.

**Fig. 6.**
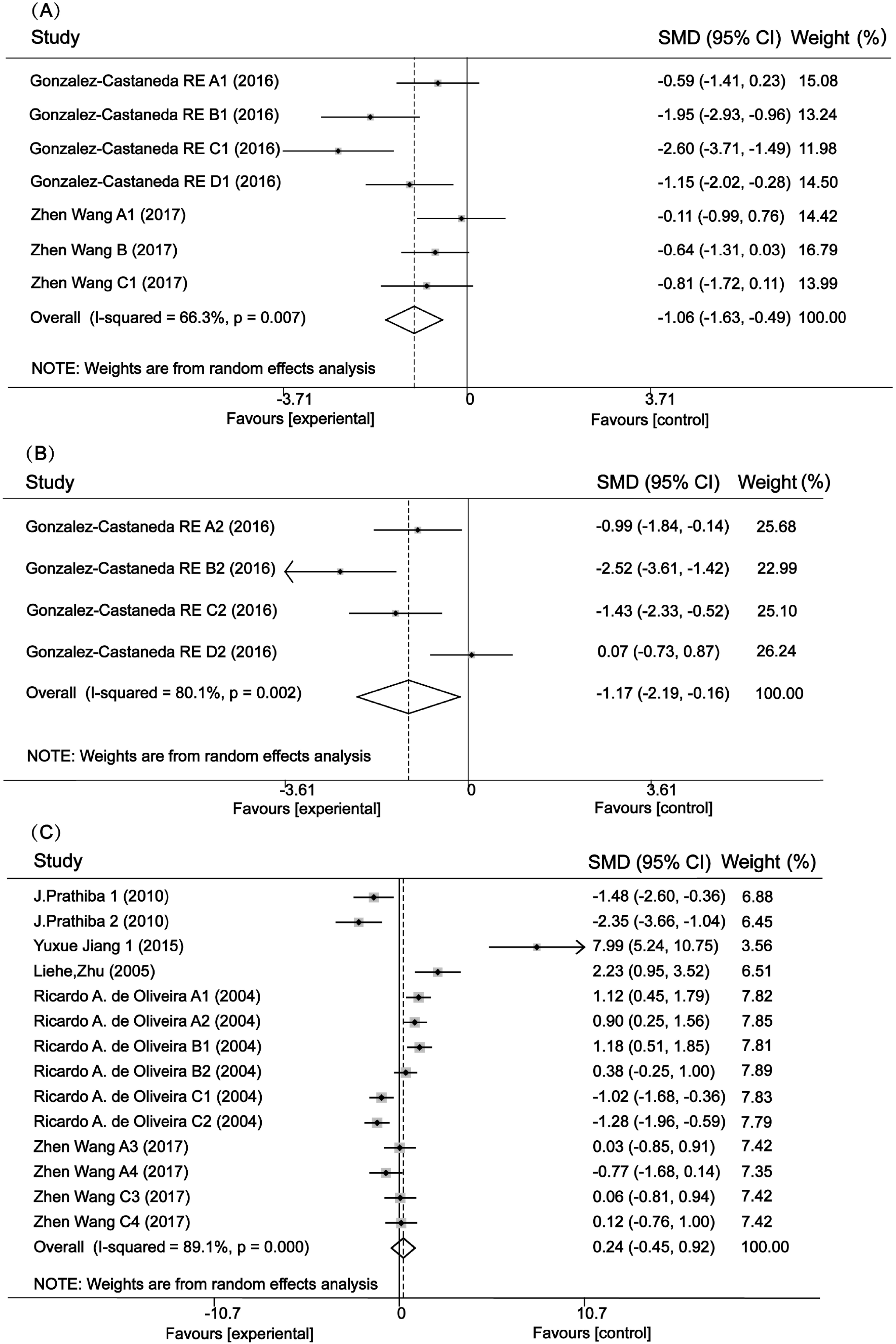
Forest plot of the random-effects model subgroup analysis of animals. A. Forest plot of the random-effects model subgroup analysis of animals regarding the sucrose consumption test. B. Forest plot of the random-effects model subgroup analysis of animals regarding the forced swimming test. C. Forest plot of the random-effects model subgroup analysis of animals regarding the open-field test.

## 4. Discussion

Sleep accounts for about one-third of human life, and it is well known that sleep maintains physical strength, restores energy, promotes growth and development, and delays aging and disease. Lack of sleep or fragmented sleep can lead to listlessness, decreased alertness, and decreased concentration at work. Severe sleep deprivation can lead to physical injury and even diseases, such as coronary heart disease, hypertension, arrhythmia, diabetes and obesity, weakened immunity, and death from overwork [27]. Moderate SD, however, may be regarded as an excellent option for an accelerated response to the treatment of depression since it is well tolerated and is devoid of the potential for drug interaction[28]. After treatment with SD, depressive symptoms have been shown to be relieved, although symptoms return to the same intensity within a few days [15].

Despite these findings, the therapeutic effect of SD is controversial. Much like the extreme of sleep deprivation, a long duration of SD may cause great harm to the human body [28]. If the duration of SD is short, the results may be affected by depression relapse [17]. When combined with other antidepressant treatments, SD may enhance its effectiveness. Other treatments for depression have included light therapy and pharmacologic treatment. One study found that light therapy was effective for seasonal depression [29], and a meta-analysis found that BLT seemed efficacious, especially when administered for 2–5 weeks and as monotherapy [10]. However, light therapy alone was not as effective as SD [30]. As for medication alone, the effects are slow to manifest, and side effects may cause patients to change medication or refuse it altogether [7, 8]. For these reasons, many researchers have tried combining antidepressants with sleep deprivation, BLT, or CBT to form integrated antidepressant treatments, which have been shown to have positive effects [31, 32]. Studies that combined SD with selective serotonin reuptake inhibitors (SSRIs) showed an antidepressant effect.

The mechanism of SD in treating depression is very complex and can be interpreted based on monoaminergic neurotransmission, neuroplasticity, and gene expression. Brain-derived neurotrophic factor (BDNF) levels have been shown to be reduced in individuals suffering from a major depressive disorder, and decreased levels were also negatively correlated with HAM-D scores. Use of SD has resulted in faster treatment response and increased BDNF levels [24]. One study found that in patients who achieved an antidepressant effect after SD, the expression of the circadian clock genes (e.g., RORA, DEC2, and PER1) increased, but in patients without such an effect, a significant decrease in the expression of these genes was found [33, 34].

All fourteen articles included used RCT models, which helped to improve the rigor and significance of our review. Judging from the total results of the patient studies, our data were not as obvious as in previous meta-analyses and were highly heterogeneous [7]. This was because we included new research and because we used digital software to address instances of incomplete data. After one paper [35] was removed through sensitivity analysis, the heterogeneity of the 7–14-day group decreased from 66.6% to 19.6%, indicating that the effect of SD on depression was related to SD duration. As the forest plot shows, a duration of less than 7 days had a small effect of worsening depression, a duration of 7–14 days had an antidepressant effect, and a duration of more than 14 days had the effect of worsening depression.

Several articles reported that depression symptoms returned immediately after SD and recovery, with some patients experiencing more severe depression than before [16]. In the **<**7 days group, SD only occurred once with a duration of < 36 hours. Therefore, it was very likely that the depression symptoms had recurred following a night of SD intervention, and that the results had a small effect of worsening depression [17]. Sleep loss, especially when chronic, can cause significant and cumulative neurobehavioral deficits and physiological changes, some of which may account for inattention, slowed working memory, reduced cognitive throughput, depressed mood, and perseveration of thought [36]. Thus, prolonged and repeated SD could worsen depression, which might account for the increased depression in groups with a duration of more than 14 days. With 7–14 days of SD, the interference of the first two conditions might be slightly avoided, thus providing a better therapeutic effect. The heterogeneity of this group mainly came from differences in sample type among the other three articles [35]. Studies have indicated that in unipolar depressed samples, the response rate to SD was 50.6%, and in samples using a mixture of unipolar and bipolar depressed patients, the response rate was 53.1% [7]. We guessed, therefore, that different types of depression samples had different response rates to SD. However, with the small amount of literature included in this study, it was impossible to clearly explore similar results.

Given the large heterogeneity in the total dataset, sources of heterogeneity were explored for potential influencing variables. The first analysis was a subgroup analysis by country. Different countries have different factors affecting the occurrence, treatment, and prognosis of depression, such as national health awareness, cultural and educational quality, medical research level, medical and social security, family economic income and social welfare, and social support systems [37, 38]. In studies from Turkey, there was an antidepressant effect of SD [-1.77 (−2.35, −1.19); I^2^=0%; P=0.586]. Meanwhile, studies from Switzerland showed that SD worsened depression [1.07 (0.51, 1.63); I^2^=0%; P=0.845]. These findings suggest that the effect of SD on depression may be related to ethnicity and nationality. Although relatively few articles were included in this study, based on the available data, we speculate that the treatment effect of SD on depression may be more likely to be observed in studies conducted in the Turkish context. An adverse effect of SD was observed in patients from Switzerland in one paper, so more studies are needed for verification. Since most patients fell into the diagnostic category of major depression, we speculated that the intervening effect of SD was more obvious for major depression. It is possible that the higher the level of depression, the more significant the therapeutic effect of SD.

In light of the above, the following points are relevant: 1) Turkey has low levels of economic and medical academic development and education, while Switzerland has high levels of those indices. 2) Two papers from Turkey included in this study [23, 24] had female patient proportions of 79% and 73.2%, while the paper from Switzerland had a female patient proportion of 39.3%. Depression levels in the Turkish studies could have been significantly related to the medical academic development. In consideration of the possible effects of racial diversity, further studies are needed to examine the effects of SD on depression across various ethnic groups.

The other subgroup analysis concerned whether there was a combination of SD with other therapies. Combined with BLT, CBT, and tricyclic antidepressive agents (TCAs), SD was shown to have no significant effect on depression. Three studies of SD combined with SSRIs showed an antidepressant effect; after removing one study [28] with fewer than seven patients in each group, which could have affected the outcome, the effect size went from [-0.58 (−1.94, 0.78)] to [-1.77 (−2.35, −1.19)], and heterogeneity went from 84.3% to 0. We suspect that heterogeneity could have been related to three aspects: 1) Patients in one of the studies were from USA, and those in the other two were from Turkey. This correlates with the results of the above analysis. 2) One study incorporated paroxetine, while sertraline was used in the other two, so heterogeneity could also have come from the use of different SSRIs. One paper’s authors stated that although sertraline and paroxetine had comparable efficacy for major depression, patients who used sertraline had a lower recurrence rate than those who used paroxetine. Sertraline was somewhat better tolerated than paroxetine and had a lower side-effect profile [39]. 3) The duration of SD in one study was less than 7 days, while it was 7–14 days in the other two studies.

In animal experiments, the overall results from data with high heterogeneity were not statistically significant and could have been related to the inconsistent effects of SD. While some studies reported antidepressant results from SD, others reported depressed results. Therefore, we used the most significant subgroup analysis (depression testing tool) to examine the sources of heterogeneity. After one paper was removed through sensitivity analysis, the heterogeneity went from 81.1% to 66.3%, and the standard mean difference (SMD) value for the sucrose consumption test changed from [-0.79 (−1.51, −0.07)] to [-1.06 (−1.63, −0.49)], which showed the effect of worsening depression. This increase in effect size could be related to the following: 1) The animals included rats and mice, which could be the source of heterogeneity. The mice and rats belonged to different species (BALB/c, C57BL, and Sprague-Dawley) with the corresponding genetic and biological characteristics. This could have led to some differences in the sensitivity of animals to SD and sucrose consumption experiments. We suspect that mouse models are more likely to derive depressive effects from SD, but further research is needed to verify this. 2) The removed paper proposed building a depression model using the chronic unpredictable stress method. Other groups did not explicitly note the establishment of the depression model. We did not consider whether an animal model for depression was clearly established as an inclusion criterion because we believed the experiments to test the degree of depression had an effect on the animal’s mental condition. However, the establishment of a depression model could cause animals to be consistent and reduce the effect of different depression experiments on the test results. Based on our sensitivity analysis, we believed the source of heterogeneity could be the depression model. In the sucrose consumption test, those without a depression model were more likely to be depressed by SD. 3) The literature that had been excluded did not mention the type of SD used, but the rest of the group used paradoxical SD. Although the background in the literature was not rich enough, we suspected that paradoxical SD was more likely to worsen depression.

Sensitivity analysis performed on the forced swimming test changed its effect size from [-0.71 (−1.53, 0.12)] to [-1.17 (−2.19, −0.16)], and heterogeneity went from 80.3% to 80.1% after one paper was excluded and reflected an antidepressant effect. Changes in effects could come about in two ways: 1) The excluded paper was from China, and the other papers were from Mexico. There is a certain gap in the level of scientific research development between China and Mexico, and there could have been differences in the experimental design, degree of rigor, scientific research concept, and environmental climate. Even if the living environment of the mice was controlled in the experimental design, it might still be affected by the local environment. We hypothesized, therefore, that between Mexico and China, perhaps the Mexican studies were more likely to show the therapeutic effect of SD on depression. 2) The animal species in the excluded paper were of the C57BL/6J type, while the others were BALB/c. Inbred C57BL/6 mice show low spontaneous activity, poor ability to explore novel environments, and proneness to behavioral despair under acute stress stimulation [40]. Therefore, in the forced swimming experiment, the expression of depression in C57BL/6J mice could have been influenced by their biological characteristics in that they were more likely to show depressive symptoms, and we suspect the therapeutic effect of SD is more pronounced in BALB/c mice. The study using an open field trial to test depression found that SD had no significant effect on depression and had high heterogeneity, which could have been related to complex experimental methods, including multiple unit tests and uneven quality analysis.

This meta-analysis included patient studies to explore the clinical effects of SD and to find the best way to use SD to treat depression. Because some monitoring indicators, such as changes in neurotransmitters, are more likely to show up in animals, analysis of animal studies was used to address the deficiencies encountered with patient analysis of SD treatments. Regardless of the reason, we should be aware of the risks involved in interpreting the efficacy and effectiveness of SD based on the discrepancies between rodent and human research.

In animal studies, many documents showed that SD affects depressive episodes by altering the regulation of serum corticosterone[41], BDNF[24], and other neurotransmitters[42]. Sleep deprivation is closely linked with the downregulation of miR-10B and possibly the upregulation of BDNF in the hippocampus in rats subjected to chronic unpredictable stress[43]. Sleep deprivation can decrease serum corticosterone levels of rats with depression, and SD improves depression in depressive rats, including hyperactivity of hypothalamus-pituitary-adrenal gland axis[41]. For the overall meta-analysis, no effect of SD on depression was observed in patients and animals, although significant effects were detected as a consequence of SD in the sensitivity analysis. Because there are differences between the effects of SD on animals and patients under certain conditions, some explanations can be proposed for the discrepancies. The first is the differences between species. Second, the methods used to test the effect of SD are qualitatively different. Depression is detected in animals using objective behavioral indicators, while it is measured in patients using subjective scoring scales. Preclinical animal studies that use a non-applicable method to assess a human condition may lack coherence and meaning in their results. It is imperative that researchers performing animal studies use the best methods available to acquire the best possible results. As this article shows, when studying the antidepressant effect of SD, the forced swimming trial can be used; when studying the depressive effect of SD, the sucrose consumption test can be used. In addition, BALB/c mice are also more likely to show the antidepressant effects of SD. By such selection, it is possible to study the mechanism of sleep deprivation in animals with a smaller margin of error.

### 5.1 Clinical and experimental implications

Based on this study’s findings, confining the duration of SD treatment to 7–14 days could be a clinically feasible way to enhance its therapeutic effect. Regarding combination treatment, SD and SSRI medications can be attempted. Combined with clinical practice, TCAs give more troublesome side effects and have the potential for fatal overdose, so SSRIs may be safer [44]. This study did not include papers with methods combining three or more therapies, so further study is needed. In animal research, the same trend method can be used to reduce the interference of depression evaluation tests on the results. When studying the antidepressant effect of SD, the forced swimming trial can be used; when studying the depressive effect of SD, the sucrose consumption test can be used. The above findings belong to the speculative aspects of this study, which need to be refined and improved by more extensive studies.

### 5.2 Study limitations

Aside from the above-mentioned speculations, several limitations should be noted. The quantity of literature included in this study was small, which could make the results less convincing. Second, the data derived using digital software were different from the actual study results, meaning there was a certain degree of data error. Third, there was no uniform model for depression in animals, and the differing results could have been produced by the depression tests that were employed. Future research should target such limitations.

## 6. Conclusion

Our meta-analysis showed that SD could be an effective antidepressant measure when the treatment duration is 7–14 days. Meanwhile, a duration of less than 7 days had a small effect of worsening depression, while a duration of more than 14 days had the effect of worsening depression. Additionally, in animal studies, depression was measured using the forced swimming experiment, which showed more antidepressant effects, while using the sucrose consumption test had the effect of worsening depression. These findings suggest that SD should be used as an intervention for depressed people within specific parameters. Further high-quality research with long-term follow-ups is needed to strengthen the evidence.

## List of abbreviations

BDNF: brain-derived neurotrophic factor
BLT: bright light therapy
CBT: cognitive behavioral treatment
HAMD: Hamilton depression scale
PSD: partial sleep deprivation
RCT: randomized controlled trials
rTMS: repetitive transcranial magnetic stimulation
SD: sleep deprivation
SMD: standard mean difference
SSRIs: selective serotonin reuptake inhibitors
TCAs: tricyclic antidepressive agents
TSD: total sleep deprivation

## Acknowledgments

This work was funded by the National Natural Science Foundation (nos. 81560059 and 81760058), Public Welfare Technology Applied Research Projects in Zhejiang Province (no. 2016C33191), Zhejiang Medical Health Science and Technology Project (no. 2019KY724, 2020KY332), the Scientific Research Fund of Shaoxing University (no. 20125025), and the National Training Program of Innovation and Entrepreneurship for College Students (no. 2017R10349001).

